# SARS-CoV-2 envelope protein topology in eukaryotic membranes

**DOI:** 10.1101/2020.05.27.118752

**Authors:** Gerard Duart, Ma Jesús García-Murria, Brayan Grau, José M. Acosta-Cáceres, Luis Martínez-Gil, Ismael Mingarro

## Abstract

Coronavirus E protein is a small membrane protein found in the virus envelope. Different coronavirus E proteins share striking biochemical and functional similarities, but sequence conservation is limited. In this report, we studied the E protein topology from the new SARS-CoV-2 virus both in microsomal membranes and in mammalian cells. Experimental data reveal that E protein is a single-spanning membrane protein with the N-terminus being translocated across the membrane, while the C-terminus is exposed to the cytoplasmic side (Nt_lum_/Ct_cyt_). The defined membrane protein topology of SARS-CoV-2 E protein may provide a useful framework to understand its interaction with other viral and host components and establish the basis to tackle the pathogenesis of SARS-CoV-2.

## INTRODUCTION

The coronavirus disease 19 (COVID-19), an extremely infectious human disease caused by coronavirus SARS-CoV-2, has spread around the world at an unprecedented rate, causing a worldwide pandemic. While the number of confirmed cases continues to grow rapidly, the molecular mechanisms behind the biogenesis of viral proteins are not fully unraveled. The SARS-CoV-2 genome encodes for up to 29 proteins, although some may not get expressed [1]. The viral RNA is packaged by the structural proteins to assemble viral particles at the ERGIC (ER-Golgi intermediate compartment). The four major structural proteins are the spike (S) surface glycoprotein, the membrane (M) matrix protein, the nucleocapsid (N) protein, and the envelope (E) protein. These conserved structural proteins are synthesized from sub-genomic RNAs (sgRNA) encoded close to the 3’ end of the viral genome 44 [2].

Among the four major structural proteins, the E protein is the smallest and has the lowest copy number of the membrane proteins found in the lipid envelope of mature virus particles (reviewed [3,4]). However, it is critical for pathogenesis of other human coronaviruses [5,6]. Interestingly, the sgRNA encoding E protein is one of the most abundantly expressed transcripts despite the protein being low copy number in mature viruses [1]. It encodes a 75 residues long polypeptide with a predicted molecular weight of ∼8 kDa. Two aliphatic amino acids (Leu and Val) constitute a substantial portion (36%, 27/75) of the E protein, which accounts for the high grand average of hydropathicity (GRAVY) index of the protein (1.128), as calculated using the ExPASy ProtParam tool (https://web.expasy.org/protparam/). Comparative sequence analysis of the E protein of SARS-CoV-2 and the other six known human coronaviruses, do not reveal any large homologous/identical regions (Figure 1), with only the initial methionine, Leu39, Cys40 and, Pro54 being ubiquitously conserved. With regard to overall sequence similarity SARS-CoV-2 E protein has the highest similarity to SARS-CoV (94.74%) with only minor differences (Figure 1B), and the lowest with HCOV-NL63 (18.46%). These findings are consistent with the phylogenetic tree proposed based on the amino acid sequences of the human coronavirus E proteins using ClustalW (Figure 1C).

**Figure 1.**
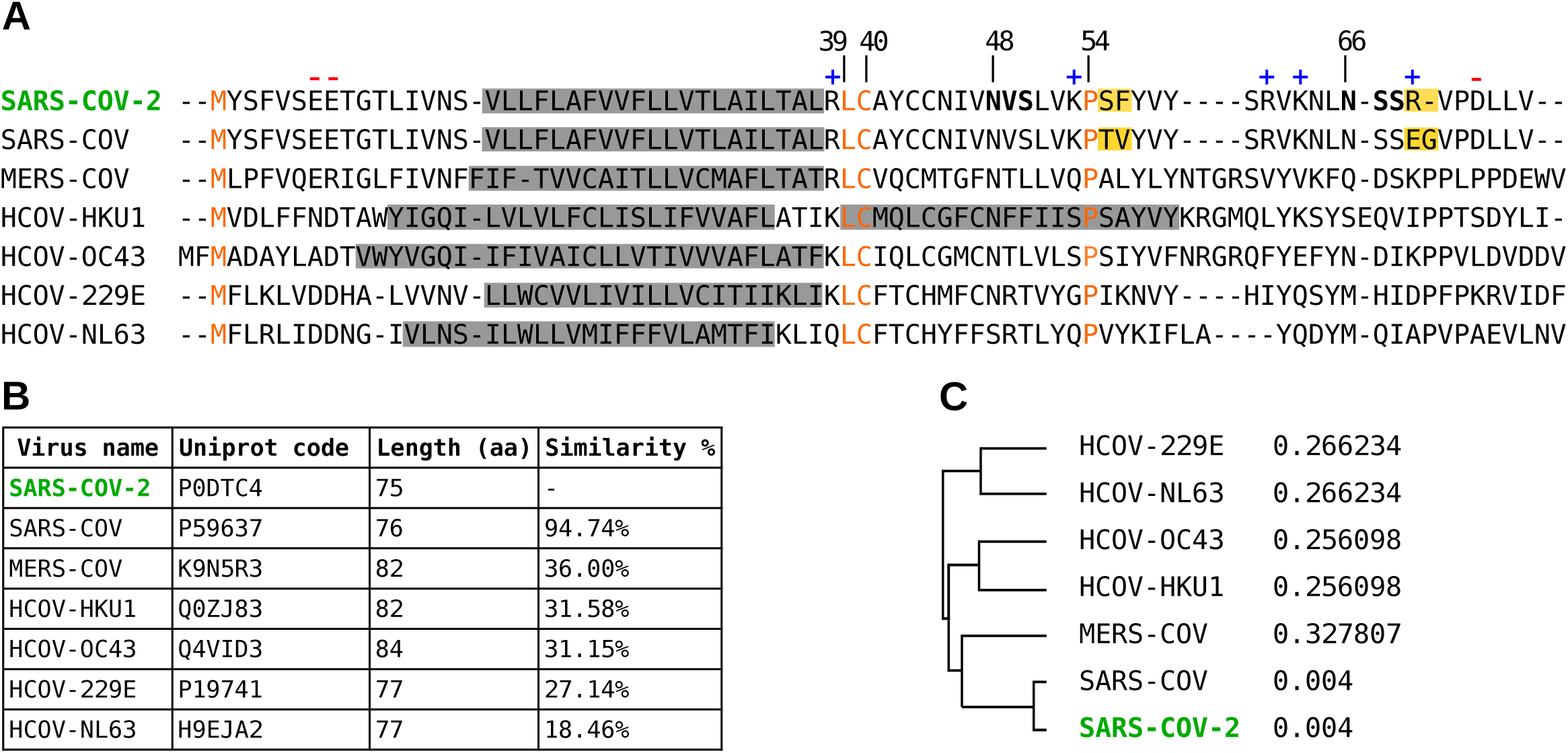
(A) Multi-alignment of amino acid sequences of the E protein of SARS-CoV-2 and the other six human coronavirus. SARS-CoV severe acute respiratory syndrome coronavirus (UniProt P59637), MERS-CoV Middle East respiratory syndrome coronavirus (UniProt K9N5R3), HCoV-HKU1 (UniProt Q0ZJ83), HCoC-OC43 (UniProt Q4VID3), HCoC-229E (UniProt P19741), HCoV-NL63 (UniProt Q5SBN7). Predicted TM segments at UniProt are highlighted in a grey box. Native predicted glycosylation acceptor sites in SARS-CoV-2 are shown in bold and charged residues highlighted with + or – symbols on top. Conserved residues are shown in orange. Differences between SARS-CoV-2 and SARS-CoV are highlighted as yellow boxes. (B) Phylogenetic data and (C) tree obtained with Clustal Omega (EMBL-EBI) using the default parameters.

## RESULTS AND DISCUSSION

Computer-assisted analysis of the SARS-CoV-2 E protein amino acid sequence using seven popular prediction methods showed that all membrane protein prediction algorithms except MEMSAT-SVM suggested the presence of one transmembrane (TM) segment located roughly around amino acids 12 to 39 (Table 1), which is not predicted as a cleavable signal sequence according to SignalP-5.0 [7]. Regarding E protein topology, TMHMM and Phobius predicted an N-terminus cytosolic orientation, whilst MEMSAT-SVM, TMpred, HMMTop and TOPCONS predicted an N-terminus luminal orientation. Firstly, we performed *in vitro* E protein transcription/translation experiments in the presence of ER-derived microsomes and [_35_S]-labeled amino acids. The membrane insertion orientation of the predicted TM segment into microsomal membranes was based on N-linked glycosylation and summarized in Figure 2 (top).

**Table 1.**
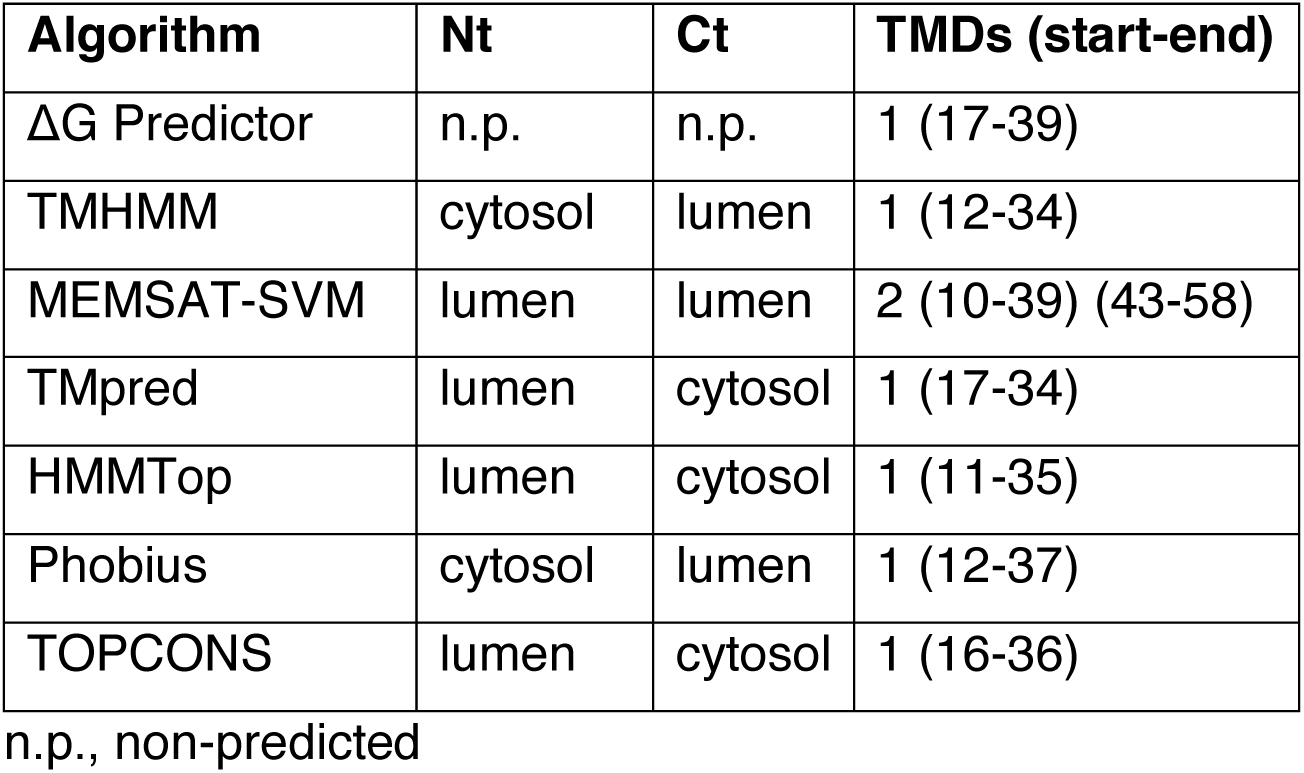
Computer analysis of the SARS-CoV-2 E protein amino acid sequence topology

**Figure 2.**
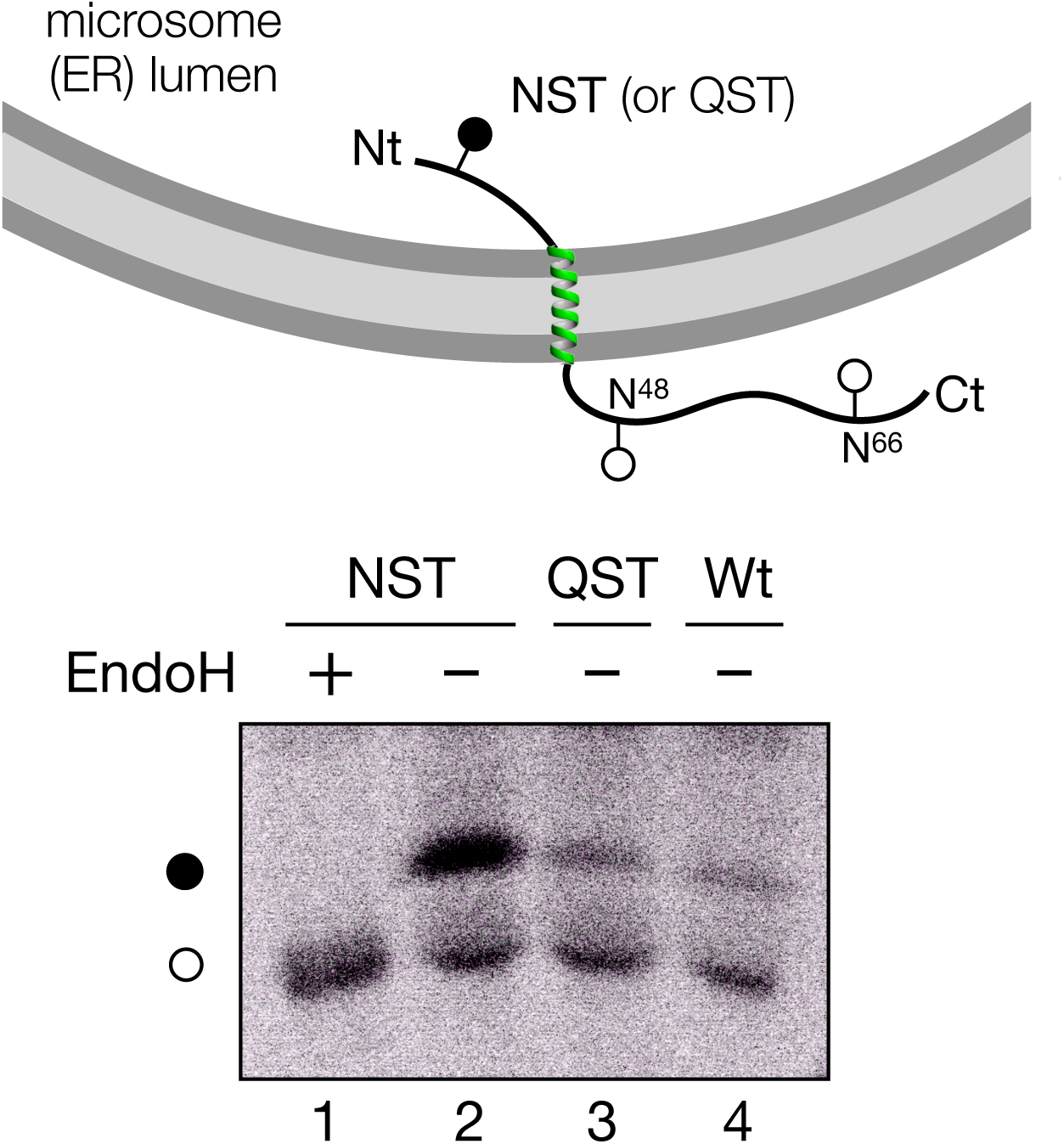
Translocon-mediated insertion of E protein variants into microsomal membranes. (Top) Schematic representation of E protein constructs. (Bottom) *In vitro* translation in the presence of microsomes of the different E protein constructs. Construct containing inserted asparagine and threonine residues at positions 3 and 5 (NST; lanes 1-2) or glutamine and threonine at positions 3 and 5 (lane 3), and wild-type variants (lane 4) were translated in the presence of microsomes. NST variant was split and half of the sample was Endo H treated (lane 1). Bands of non-glycosylated and glycosylated proteins are indicated by white and black dots, respectively. The gel is representative of at least four independent experiments.

N-linked glycosylation has been extensively used as topological reporter for more than two decades [8]. In eukaryotic cells, proteins can only be glycosylated in the lumen of the ER because the active site of oligosaccharyl transferase, a translocon-associated protein responsible for N glycosylation [9], is located there [10]; no N-linked glycosylation occurs within the membrane or in the cytosol. It is important to note that two possible N-linked glycosylation sites are located C-terminally of the predicted TM segment in E protein wild-type sequence at positions N48 and N66 (Figure 1). However, N48 is not expected to be modified even if situated lumenally due to the close proximity of this glycosylation acceptor site to the membrane if the hydrophobic region is recognized as TM by the translocon [11,12]. Thus, mono-glycosylation (at N66) would serves as a C-terminal translocation reporter. To test N-terminal translocation a construct was engineered where a predicted highly efficient glycosylation acceptor site (NST) was designed at the N-terminus. When E protein constructs were translated *in vitro* in the presence of microsomes the protein was significantly glycosylated when the N-terminal designed glycosylation site was present, as shown by the increase in the electrophoretic mobility of the slower radioactive band after an endoglycosidase H (Endo H) treatment (Figure 2, lanes 1 and 2). However, when a control (QST) that is not a glycosylation acceptor site (lane 3) or the wild-type (lane 4) sequences were translated, E protein molecules were minimally glycosylated. Since multiple topologies have been reported for previous coronavirus E proteins [13-17], SARS-CoV-2 E protein insertion into the microsomal membranes in two opposite orientations cannot be discarded, but being dominant an Nt_lum_/Ct_cyt_ orientation.

To analyse protein topology in mammalian cells, a series of E protein variants tagged with c-myc epitope at the C-terminus were transfected into HEK-293T cells. As shown in Figure 3A, only an E protein construct harbouring the N-terminal engineered acceptor site was efficiently modified (lanes 1-4), denoting an N-terminal ER luminal localisation (Nt_lum_). Several topological parameters have been proposed to govern membrane protein topology, among them the preferential distribution of positively charged residues in the cytosol (‘positive-inside rule’) has been established as the primary topology determinant both experimentally [18] and statistically [19]. E protein is a single-spanning membrane protein with an even net charge distribution on both sides of the membrane. There are only eight charged residues along the protein sequence, two negatively charged residues preceding the TM segment and five positively and one negatively charged residues at the C-terminal domain (Fig. 1A), that nicely correlates the observed topology with the ‘positive-inside rule’. However, negatively charged residues have also been proved to significantly affect the topology [20]. To test the robustness of the observed topology, we added an optimized Ct glycosylation tag [21] and replaced the two negatively charged residues located in the translocated N-terminal domain (E7 and E8) by two lysine residues (Fig. 3B). In cells expressing this mutant E protein (EE>KK), the protein retained its C-terminal tail at the cytosolic side of the membrane as indicated by the absence of glycosylated forms (Fig. 3B, lanes 3 and 4). These data reveal that topological determinants have only a minor effect on viral membrane protein topology as previously demonstrated for other viruses [22], and suggest that viral membrane protein topology could have co-evolved with the protein environment of its natural host, ensuring proper membrane protein orientation. Altogether, the present *in vivo* results demonstrated that SARS-CoV-2 E protein is a single-spanning membrane protein with an Nt_lum_/Ct_cyt_ orientation in mammalian cell membranes. Similarly, SARS-CoV E protein was shown to mainly adopt an Nt_lum_/Ct_cyt_ topology in infected and transiently expressing mammalian cells [23]. This topology is compatible with the ion channel capacity described previously [24], and with the recently published pentameric structural model of SARS-CoV E protein in micelles [25], in which the C-terminal tail of the protein is α-helical and extramembrane.

**Figure 3.**
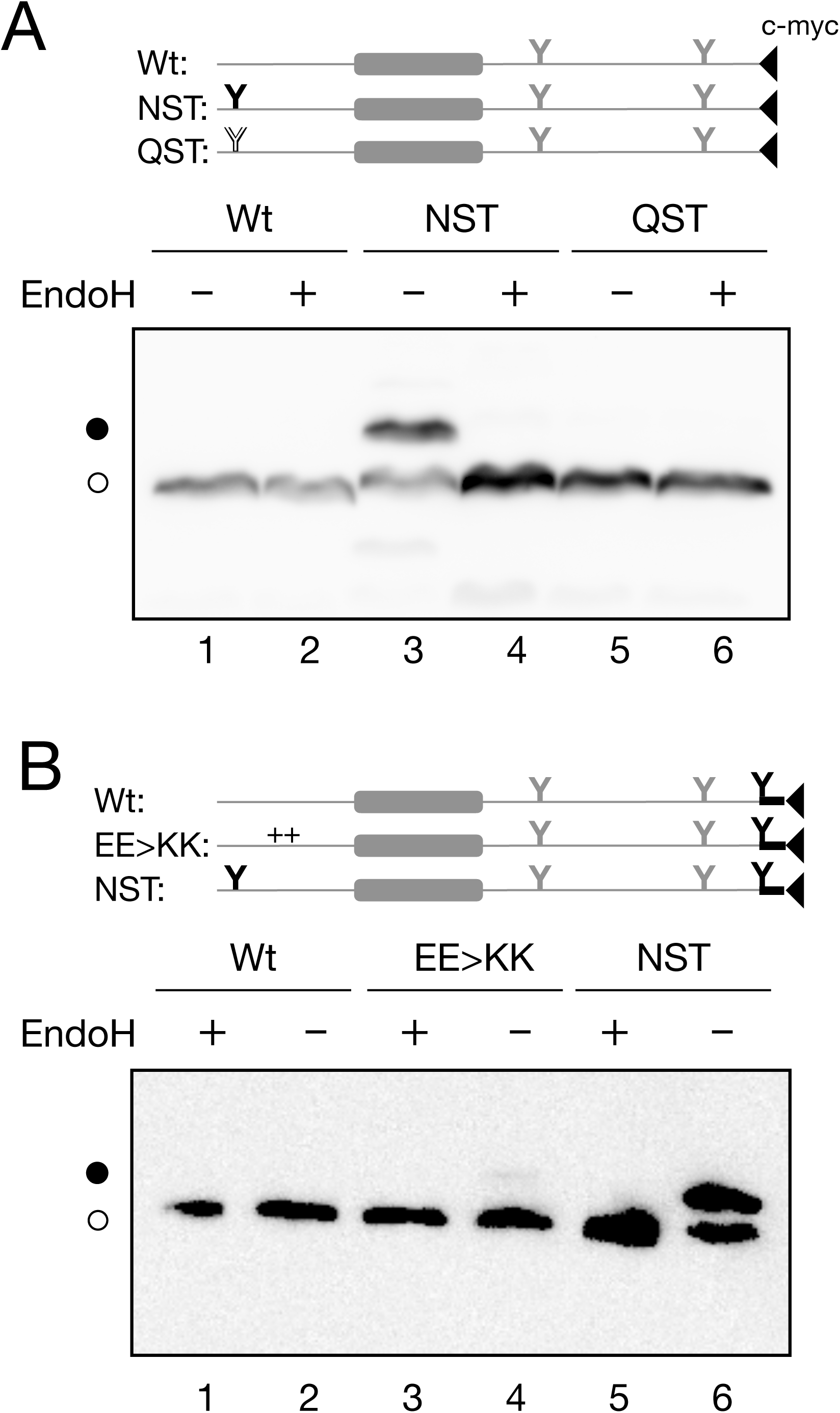
E protein topology in mammalian cells. To determine the topology *in vivo* HEK-293T cells were transfected with C-terminal tagged (c-myc) E protein variants. (A) Constructs encoding wild-type (Wt; lanes 1 and 2), inserted asparagine and threonine at positions 3 and 5 (NST; lanes 3 and 4) or glutamine and threonine at positions 3 and 5 (QST; lanes 5 and 6) were Endo H (+) or mock (-) treated. Filled and empty Y-shaped symbols denoted acceptor (NST) and non-acceptor (QST) glycosylation sites, respectively. (B) Additionally, we included constructs containing similar Wt (lanes 1 and 2), replaced glutamic acids at positions 7 and 8 by lysine residues (EE>KK; lanes 3 and 4) or NST (lanes 5 and 6) variants with an extra glycosylation site inserted at the Ct end of the protein. Once again, to confirm the glycosylated nature of the higher molecular weight bands, samples were either Endo H (+) or mock (-) treated. Designed glycosylation sites and tags are shown in black, while native E protein features are shown in gray.

The membrane topology described here, would allow the cytoplasmic C-terminal tail of the E protein to interact with the C-termini of M and/or S SARS-CoV-2 membrane embedded proteins [3], and/or with Golgi scaffold proteins as previously described for other coronaviruses [26], to induce virus budding or influence vesicular traffic through the Golgi complex by collecting viral membrane proteins for assembly at Golgi membranes. Future experiments will have to unravel whether these functions involve the SARS-CoV-2 E-protein.

## EXPERIMENTAL METHODS

### Enzymes and chemicals

TNT T7 Quick for PCR DNA was from Promega (Madison, WI, USA). Dog pancreas ER rough microsomes were from tRNA Probes (College Station, TX, USA). EasyTag™ EXPRESS35S Protein Labeling Mix, [_35_S]-L-methionine and [_35_S]-L-cysteine, for *in vitro* labeling was purchased from Perkin Elmer (Waltham, MA, USA). Restriction enzymes were from New England Biolabs (Massachusetts, USA) and endoglycosidase H was from Roche Molecular Biochemicals (Basel, Switzerland). PCR and plasmid purification kits were from Thermo Fisher Scientific (Ulm, Germany). All oligonucleotides were purchased from Macrogen (Seoul, South Korea).

### Computer-assisted analysis of E protein sequence

Prediction of transmembrane segments was done using up to 7 of the most common methods available on the Internet: ΔG Predictor [27,28] (http://dgpred.cbr.su.se/), TMHMM [29] (http://www.cbs.dtu.dk/services/TMHMM/), MEMSAT-SVM [30] (http://bioinf.cs.ucl.ac.uk/psipred/), TMpred (https://embnet.vital-it.ch/software/TMPRED_form.html), HMMTop [31] (http://www.enzim.hu/hmmtop/), Phobius [32] (http://phobius.sbc.su.se/) and TOPCONS [33] (http://topcons.net/). All user-adjustable parameters were left at their default values.

### DNA Manipulation

Full-length E protein was synthesized by Invitrogen (*GeneArt* gene synthesis) and subcloned into *Kpn*I linearized pCAGGS using In-Fusion HD cloning Kit (Takara) according to the manufacturer’s instructions. For *in vitro* assays, DNA was amplified by PCR adding the T7 promoter and the relevant glycosylation sites during the process. N-terminal NST glycosylation site was designed by inserting an asparagine and a threonine before and after Ser3, respectively. Control no-glycosylable QST site was introduced in similarly inserting a glutamine residue instead of an asparagine. All E protein variants were obtained by site-directed mutagenesis using QuikChange kit (Stratagene, La Jolla, California) and were confirmed by sequencing the plasmid DNA at Macrogen Company (Seoul, South Korea)

### Translocon-mediated insertion into microsomal membranes

E protein variants, PCR amplified from pCAGGS, were transcribed and translated using the TNT T7 Quick for PCR DNA Coupled Transcription/Translation System (Promega, USA). The reactions contained 10 *µ*L of TNT, 2 *µ*L of PCR product, 1 *µ*L of EasyTag (5 *µ*C_i_), and 0.6 *µ*L of column-washed microsomes (tRNA Probes, USA) and were incubated for 60 min at 30 °C. Translation products were ultracentrifuged (100,000 *g* for 15 min) on a 0.5 M sucrose cushion, and analyzed by SDS-PAGE. For the endoglycosidase H (Endo H) the treatment was done as previously described [34]. Briefly, the translation mixture was diluted in 120 *µ*L of PBS and centrifuged on a 0.5 M sucrose cushion (100 000 × g 15 min 4 °C). The pellet was then suspended in 50 μL of sodium citrate buffer with 0.5% SDS and 1% β-mercaptoethanol, boiled 5 min, and incubated 1 h at 37 °C with 1 unit of Endo H. Then, the samples were analyzed by SDS-PAGE.

### E protein expression in mammalian cells

E protein sequence variants were tagged with an optimized C-terminal glycosylation site [21,35] plus a c-myc epitope at their C-terminus and inserted in a pCAGGS-ampicillin plasmid. Once the sequence was verified, plasmids were transfected into HEK293-T cells using Lipofectamine 2000 (Life Technologies) according to the manufacturer’s protocol. Approximately 24 h post-transfection cells were harvested and washed with PBS buffer. After a short centrifugation (1000 rpm for 5 min on a table-top centrifuge) cells were lysed by adding 100 μL of lysis buffer (30 mM Tris-HCl, 150 mM NaCl, 0.5% Nonidet P-40) were sonicated in an ice bath in a bioruptor (Diagenode) during 10min and centrifugated. Total protein was quantified and equal amounts of protein submitted to Endo H treatment or mock-treated, followed by SDS-PAGE analysis and transferred into a PVDF transfer membrane (ThermoFisher Scientific). Protein glycosylation status was analysed by Western Blot using an anti-c-myc antibody (Sigma), anti-rabbit IgG-peroxidase conjugated (Sigma), and with ECL developing reagent (GE Healthcare). Chemiluminescence was visualized using an ImageQuantTM LAS 4000mini Biomolecular Imager (GE Healthcare).

## Acknowledgments

We thank Prof. Paul Whitley (University of Bath) for critical reading of the manuscript. This work was supported by grants PROMETEU/2019/065 from Generalitat Valenciana and COV20/01265 from ISCIII (to I.M.). G.D. was recipient of a predoctoral contract from the Spanish Ministry of Education (FPU). J.A.-C. was recipient of a predoctoral fellowship from the Ministerio de Educación (Perú). B.G. was recipient of a predoctoral contract from the University of Valencia (Atracció de Talent).

## Notes

### Competing Interest Statement

The authors have declared no competing interest.

